# lncRNA Mediated Hijacking of T-cell Hypoxia Response Pathway by *Mycobacterium tuberculosis* Predicts Latent to Active Progression in Humans

**DOI:** 10.1101/2020.04.11.037176

**Authors:** Jyotsana Mehra, Vikram Kumar, Priyansh Srivastava, Tavpritesh Sethi

## Abstract

Cytosolic functions of Long non-coding RNAs including mRNA translation masking and sponging are major regulators of biological pathways. Formation of T cell-bounded hypoxic granuloma is a host immune defence for containing infected Mtb-macrophages. Our study exploits the mechanistic pathway of Mtb-induced HIF1A silencing by the antisense lncRNA-HIF1A-AS2 in T cells. Computational analysis of in-vitro T-cell stimulation assays in progressors (n=119) versus non-progressor (n=221) tuberculosis patients revealed the role of lncRNA mediated disruption of hypoxia adaptation pathways in progressors. We found 291 upregulated and 227 downregulated DE lncRNAs that were correlated at mRNA level with HIF1A and HILPDA which are major players in hypoxia response. We also report novel lncRNA-AC010655 (AC010655.4 and AC010655.2) in cis with HILPDA, both of which contain binding sites for the BARX2 transcription factor, thus indicating a mechanistic role. Detailed comparison of infection with antigenic stimulation showed a non-random enrichment of lncRNAs in the cytoplasmic fraction of the cell in TB progressors. The lack of this pattern in non-progressors replicates indicates the hijacking of the lncRNA dynamics by Mtb. The in-vitro manifestation of this response in the absence of granuloma indicates pre-programmed host-pathogen interaction between T-cells and Mtb regulated through lncRNAs, thus tipping this balance towards progression or containment of Mtb. Finally, we trained multiple machine learning classifiers for reliable prediction of latent to the active progression of patients, yielding a model to guide aggressive treatment.

## Introduction

*Mycobacterium tuberculosis* (Mtb) is a non-motile, rod-shaped bacillus which was discovered by Robert Kock in the late 18th century. The bacterium is distantly related to Actinomycetes [1]. Dimensionally the average length of bacillus is equal to 2 microns, with 0.5 microns in width. Mtb is neither gram-positive or gram-negative rather it is stained through acid-fast staining [2]. The most widely used method for its acid-fast staining is Ziehl–Neelsen staining [2]. Mtb first infects the Macrophages through vesicle trafficking events by disrupting the antigen processing and presentation pathway which includes phagosome-lysosome fusion [3]. Once the soluble antigens are presented to the CD4+ T cells by the un-infected dendritic cells, the adaptive immune response is stimulated. As a result of which these virgin T cells become polyfunctional through antigen-driven differentiation [4]. Among all the antigens, diagnostic antigens ESAT and Ag85 play a vital role in disease progression and pathogenesis [5,6]. ESAT-6 inhibits NF-κB activation that restricts myeloid differentiation, whereas the Ag85 prevents the formation of phagolysosome [5,6]. Hypoxia-induced dormancy of Mtb is one of the primary goals of the granuloma formation by T cells and other immune cells [7]. Adaptive survival of Mtb in hypoxic granulomas has been extensively studied [8]. CD8+ T cells are the major killers of dormant Mtb as they go deep in the granulomas [9]. To withstand the hypoxic environment of granulomas, T cells induce HIF-1A which is one of the pioneer genes involved in the hypoxic-homeostasis [10].

Long non-coding RNAs (lncRNAs) are defined as transcripts of lengths exceeding 200 nucleotides that are not translated into protein and form the major part of the non-coding transcriptome. Genome-wide association studies (GWAS) have evaluated their role in disease progression and development. They play a crucial role in gene expression by controlling the translational freedom through masking and silencing events [11]. Previous findings have escalated the role of Nuclear-enriched abundant transcript 1 (lncRNA-NEAT1) as an important immune regulator of Tuberculosis (TB) prognosis [12]. 2016’s Microarray studies of Mtb infected macrophages eluted out MIR3945HG V1 and MIR3945HG V2, as novel biomarkers for TB that play a vital role in Mtb-macrophage interaction [13]. In the successive year lncRNAs role in the regulation of alpha-beta T cell activation and the T cell receptor signalling pathway have been actively studied, making them a potent early-diagnosable biomarker of TB [14]. Experimental studies also documented the effects of lncPCED1B-AS on macrophage apoptosis [15].

The present study found the mRNA silencing potential of lncRNA-HIF1A-AS2 in T cells which disrupts the hypoxia adaptation pathways in TB progressors (patients who develop active TB from latent stages). According to our proposed hypotheses, lncRNA-HIF1A-AS2 silences its anti-sense mRNA-HIF-1A and is induced by Mtb during latent to active TB transition. The adaptation mechanism is also assisted with HILPDA which is responsible for lipid accumulation as in low oxygen environments. We applied statistical procedures to find out differentially expressed genes in Mtb-infected T cells samples against Antigen-stimulated T cells samples, to decipher the Mtb T-cell interaction during active TB transition. Statistical inferences from our study show 291 upregulated lncRNAs that were correlated at mRNA level with HIF1A and HILPDA are major players in hypoxia response. Computational analysis of in-vitro T-cell stimulation assays in progressors versus latent tuberculosis patients also shows major differences in the chromosomal distribution of genes and sub-cellular localization of the DE lncRNAs.

## Methods

### 2.1 Data Collection and Curation

The data for analysis was taken from the expression profiling study carried on 150 adolescents (12-18 years), with a gap cycle of 6 months [16]. The reads were recorded using Illumina HiSeq 2000 (*Homo sapiens*) (GPL11154) and are available at Gene Expression Omnibus (GEO) with Accession ID as “GSE103147” [17]. 106 blood samples of non-progressors (Mtb infected controls) and 44 blood samples of progressors (developed TB during two years of followup) constituted the initial data set. A total of 1650 raw RNASeq reads of 150 blood samples were systematically retrieved from the GEO database through GEOquery using Bioconductor (R.3.6) [17-19]. We proceeded with 340 T-cell replicate samples which included 119 progressors replicates and 221 non-progressors replicates collected on the first day of study. 340 replicate sample data was distributed into four classes (“Mtb Infected”, “Ag85 Stimulated”, “ESAT Stimulated” and “Unstimulated”) based on the method of stimulation.

### 2.2 Construction of alignment maps

All spliced reads in FASTQ files after retrieval were aligned with the human genome (GRCh37-hg19) to produce the alignment maps [20]. Hierarchical Indexing for Spliced Alignment of Transcript (HISAT) was used for the reference-based assembly of the reads to the GRCh37-hg19 [20,21]. All the alignment maps (SAM files) were converted into binary aligned maps (BAM files) using Sam tools [22] (Fig. 1).

**Figure 1.**
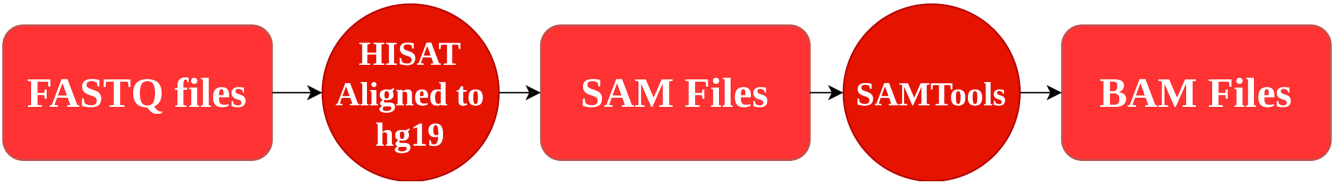
(Pipeline used for reference-based genome assembly)

### 2.3 Extraction of lncRNAs and mRNAs

Ultrafast Comprehensive Long Non-Coding RNA (UClncR) pipeline was implemented on the BAM files for extraction of the expression counts of lncRNAs [23]. 68,000 lncRNAs were generated by the UClncR pipeline [23]. After dropping 54,837 un-annotated and 2,382 insignificant (0 expressions across all replicates) lncRNAs, 10,781 lncRNAs were taken for differential expression analysis (Fig. 2(a)). Similarly, 55,190 mRNAs were generated using the HtSeq-count [24]. After filtering protein-coding mRNAs using RefSeq database, and dropping insignificant mRNAs, 16,969 mRNAs were taken for differential gene expression analysis [23, 25] (Fig. 2(b)).

**Figure 2.**
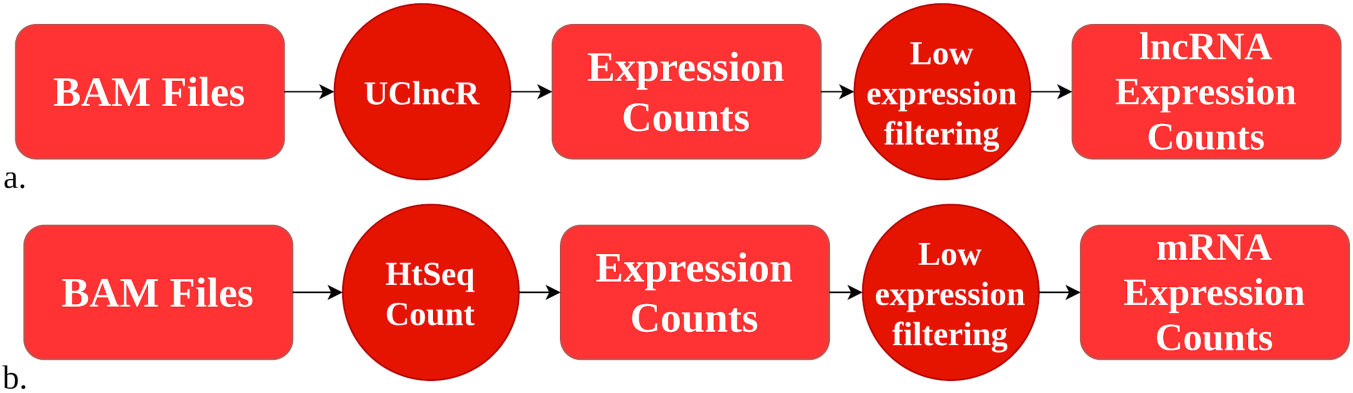
(Pipelines followed for extraction of expression counts of protein-coding mRNAs and lncRNAs; (a) Pipeline for lncRNA, filtering removed novel transcripts and low expression count across all samples; (b) Pipeline for protein-coding mRNA, filtering removed non-coding mRNA species and low expression count across all samples)

### 2.4 Exploratory Data Analysis

We plotted the expression counts of the samples class-wise (i.e. “Mtb Infected”, “Ag85 Stimulated”, “ESAT Stimulated” and “Unstimulated”) by Uniform Manifold Approximation and Projection (UMAP) [26]. Interestingly, we found the cluster among three classes (i.e. “Ag85 Stimulated”, “ESAT Stimulated” and “Unstimulated”), whereas, the “Mtb Infected” class formed an isolated cluster (Fig. 3). Therefore conducted differential gene expression analysis both class-wise (“Mtb Infected” vs “Antigen ESAT/Ag85 Stimulated”) and sample-wise (“Progressor” vs “Non-Progressor”). We dropped the “Unstimulated” class from our class-wise differential analysis as part of sub-experimental control.

**Figure 3.**
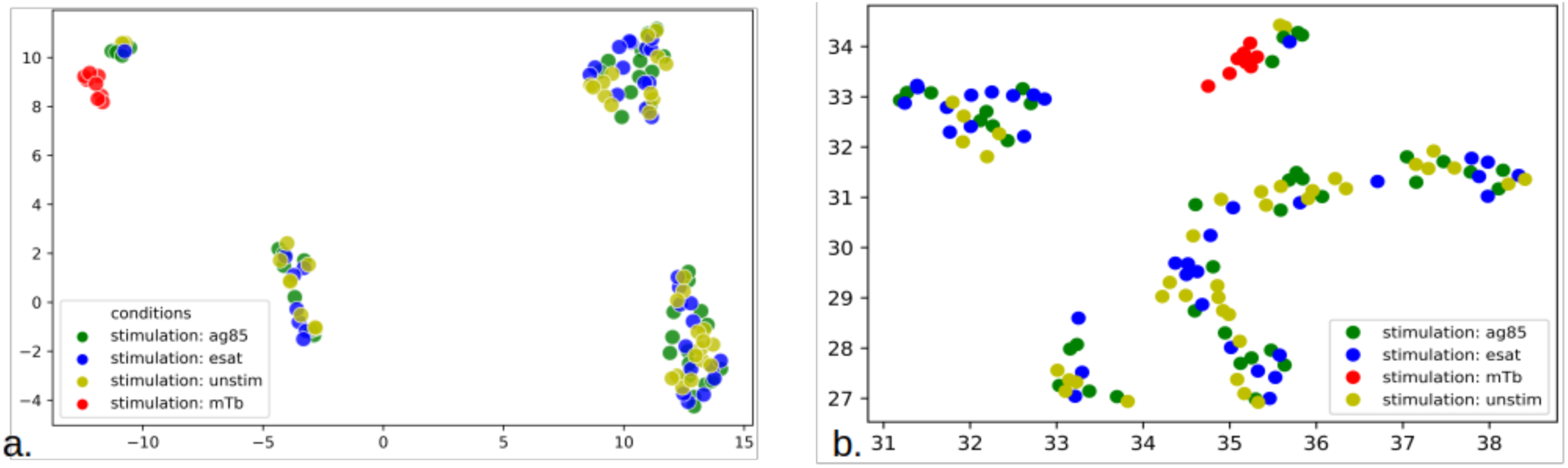
(a). Uniform Manifold Approximation and Projection for lncRNA expression counts; Classes include Stimulated by Ag85 in red, Stimulated by ESAT in green, Infected by Mtb in violet, Unstimulated in blue; Clusters of Mtb are isolated against Stimulated sets of ESAT and Ag85; (b) Uniform Manifold Approximation and Projection for mRNA expression counts; Classes include Stimulated by Ag85 in red, Stimulated by ESAT in green, Infected by Mtb in violet, Unstimulated in blue; Clusters of Mtb are isolated against Stimulated sets of ESAT and Ag85;

### 2.5 Differential Expression Analysis

Calculation of DE lncRNA and DE mRNA was done both class-wise (“Mtb Infected” vs “Antigen ESAT/Ag85 Stimulated”) and sample-wise (“Progressor” vs “Non-Progressor”) (Fig. 4). ANOVA was applied across all the samples (both class wise and sample wise) for calculation of differentially expressed elements [27]. For higher confidence false discovery rate (FDR) cut-off of <0.01 was applied. For extraction of DE lncRNA, lncDiff pipeline was implemented [28]. Whereas, for the extraction of DE mRNA, we used the EdgeR pipeline [29]. We estimated the expression status of differential elements by finding the log of two-fold changes. Differential elements (i.e. DE lncRNAs and DE mRNAs) having log2Fc < −1 were classified as downregulated elements and those having log2Fc > +1 were classified as upregulated elements. Elements with −1 < log2Fc < +1 were labeled as insignificant elements.

**Figure 4.**
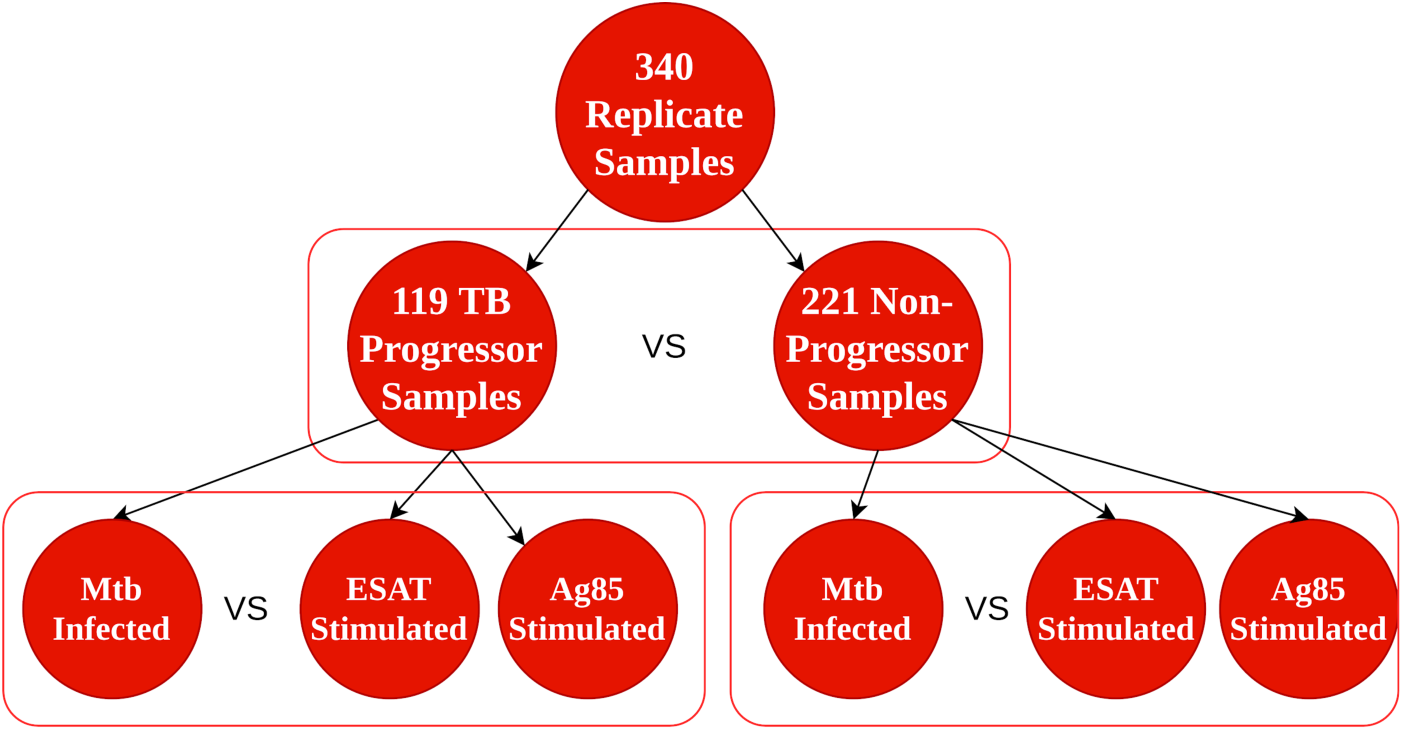
(Graphical summarization of the differential expression analysis done on the lncRNA and mRNA expression counts. Sample wise (Progressor vs Non-Progressor) and class-wise (Mtb Infected vs ESAT/Ag85 stimulated) was done independently of each other)

### 2.6 Gene Set Enrichment Analysis

As lncRNAs cannot be enriched using traditional Gene Set Enrichment Analysis (GSEA) approaches, therefore, we enriched correlated DE mRNA in place of them. The Pearson-correlation test was conducted between DE mRNA and DE lncRNA generated from the class-wise differential analysis (“Mtb Infected” vs “Antigen ESAT/Ag85 Stimulated”). The DE mRNAs from highly correlated DE lncRNA-DE mRNA pairs were used for GSEA. ShinyGO server was used to perform GSEA of the correlated mRNA [30]. For gene ontology prediction we used Gene Ontology (GO) database, and for pathway enrichment, we used Kyoto Encyclopedia of Gene and Genome (KEGG) database via ShinyGO server [30-32]. The most significant GO terms and pathways were filtered using the p-value cut-off of <0.05.

### 2.7 Mechanistic Analysis

Biomart service was used to map the genomic features such as chromosomal locations, HGNC symbols and descriptions of the correlated DE lncRNA-DE mRNA pairs [33]. The correlated pairs with the heterogeneous chromosomal numbers were programmatically dropped. We further filtered the data set based on the start base-pair positions of the differential elements in the correlated DE lncRNA-DE mRNA pairs. Pairs having a distance of less than 1000 bp were selected for further analysis.

### 2.8 Transcription factor binding and Sub-cellular localization

The JASPER database was used to retrieve the profiles of the transcription factor binding sites in the transcripts of *Homo sapiens* [34]. To map each transcription site to their respective chromosome location we used Ciiider with GRCh37.p12 assembly [20,35] For subcellular localization analysis, we specifically used the data of B-cells from LncATLAS database and programmatically mapped them to the respective DE lncRNAs (class-wise) [36]. For deriving the coding potential of nuclear elements we used Coding Potential Calculator [37].

### 2.9 Progression Classifier Construction

Since the DE lncRNA (class-wise in progressors) profiles showed exclusive enrichment in the cytosol, we leveraged the signal by combining the DE lncRNA enriched in the cytoplasm (class-wise in progressors) with DE mRNAs (sample-wise). The dataset was divided into training and testing sets with 80:20 ratio. We implemented the Random Forest (RF), Logistic Regression (LR), Support Vector Machine (SVM) and Decision Tree (DT) classifiers to check the accuracy, sensitivity and specificity of the model. Hyperparameters such as number of iterations (in SVM, LR; max_iter = 10000), random state (as 12345), cost function (in SVM, LR; C = 1000).

## Results

### 3.1 Differential Expression analysis

529 DE lncRNA were derived from the class-wise testing in progressors (i.e. Mtb Infected vs ESAT/Ag85 stimulated) out of which 291 lncRNA were upregulated and 227 lncRNA were downregulated in Mtb infected progressor replicates (Fig. 5(a)). As for class-wise testing in non-progressors out of 1,596 DE lncRNA, 808 lncRNA were upregulated and 490 lncRNA were downregulated (Fig. 5(c)). Similarly, 2506 DE protein-coding mRNA were derived from class-wise testing in progressors out of which 355 protein-coding mRNA were upregulated and 1210 protein-coding mRNA were downregulated in Mtb infected progressor replicates (Fig. 5(b)). For class-wise testing in non-progressors out of 5,860 DE protein-coding mRNA, 1021 protein-coding mRNA were upregulated and 882 protein-coding mRNA were downregulated (Fig. 5(d)). In the case of sample-wise testing (Progressor vs Non-Progressors), 4,765 DE protein coding mRNA were derived from sample-wise testing, out of which 3,752 protein-coding mRNA were upregulated and 3 protein-coding mRNA were downregulated.

**Figure 5.**
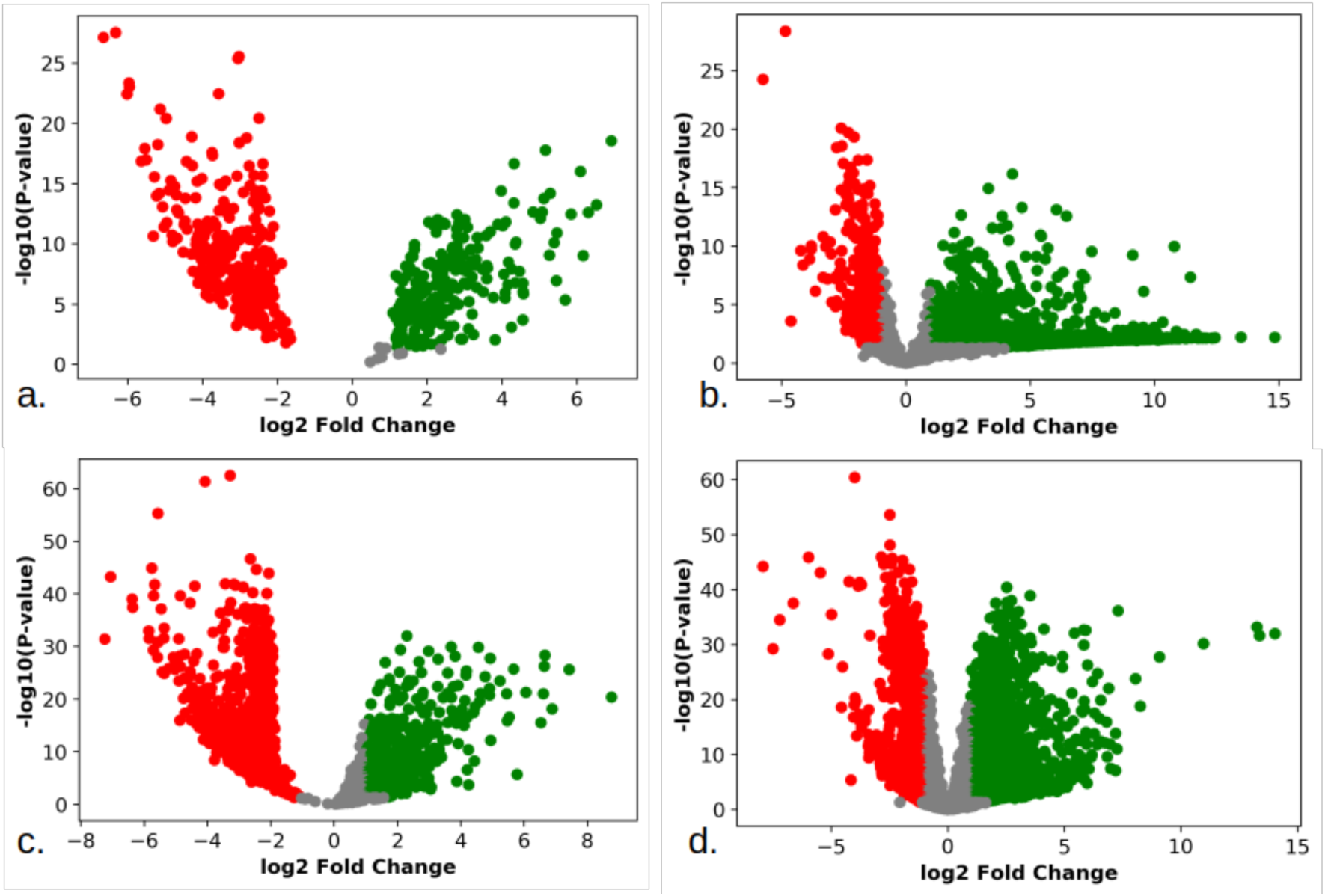
(Volcano plots differential elements (DE lncRNA and DE protein-coding mRNA) derived from class-wise testing (i.e. “Mtb Infected” vs “Ag85/ESAT stimulated”); Upregulated elements are shown in green, downregulated elements are shown in red and insignificant elements are shown in grey; (a) Volcano plot for DE lncRNA in class-wise testing of TB progressors; (b) Volcano plot for DE protein-coding mRNA in class-wise testing of TB progressors; (c) Volcano plot for DE lncRNA in class-wise testing of non-progressors; (b) Volcano plot for DE protein-coding mRNA in class-wise testing of non-progressors;)

### 3.2 GO and pathway enrichment analysis

We generated correlated pairs of DE lncRNA and DE protein-coding mRNA derived from class-wise testing for the purpose of GSEA. The correlated DE protein-coding mRNA from TB progressors class-wise testing were mainly enriched in Thermogenesis and Parkinson’s Disease via KEGG (Table 1, Fig. 6). The KEGG enrichment of correlated DE protein-coding mRNA from non-progressors class-wise testing enriched random metabolic pathways. For bias reduction and confidence, we also formed DE lncRNA - expressed mRNA (i.e. all mRNA expression counts) pairs and performed GSEA and found the same enrichment results. Close examination of the enriched pathway showed the involvement of mitochondria in thermogenesis and Parkinson’s Disease.

**Table 1.** (GSEA of correlated DE protein-coding mRNA from TB progressors class-wise testing)

| FDR | Enrichment Type | Enrichment Term | Count |
| --- | --- | --- | --- |
| 0.006757268 | KEGG | Thermogenesis | 20 |
| 0.013159226 | KEGG | Parkinson’s Disease | 14 |
| 0.015379000 | KEGG | Systemic Lupus Erythematosus | 13 |
| 0.000358735 | GO: Molecular Function | Nucleic Acid Binding | 190 |
| 0.000358735 | GO: Molecular Function | Enzyme Binding | 112 |
| 0.000840228 | GO: Molecular Function | DNA Binding | 123 |
| 0.000000001 | GO: Biological Process | Organelle Organization | 201 |
| 0.005500517 | GO: Biological Process | Regulation of Gene Expression | 198 |
| 0.005500517 | GO: Biological Process | Phosphorylation | 116 |
| 0.000000006 | GO: Cellular Compartment | Nuclear Lumen | 211 |
| 0.000000006 | GO: Cellular Compartment | Nuclear Part | 227 |
| 0.000000017 | GO: Cellular Compartment | Nucleoplasm | 184 |

**Figure 6.**
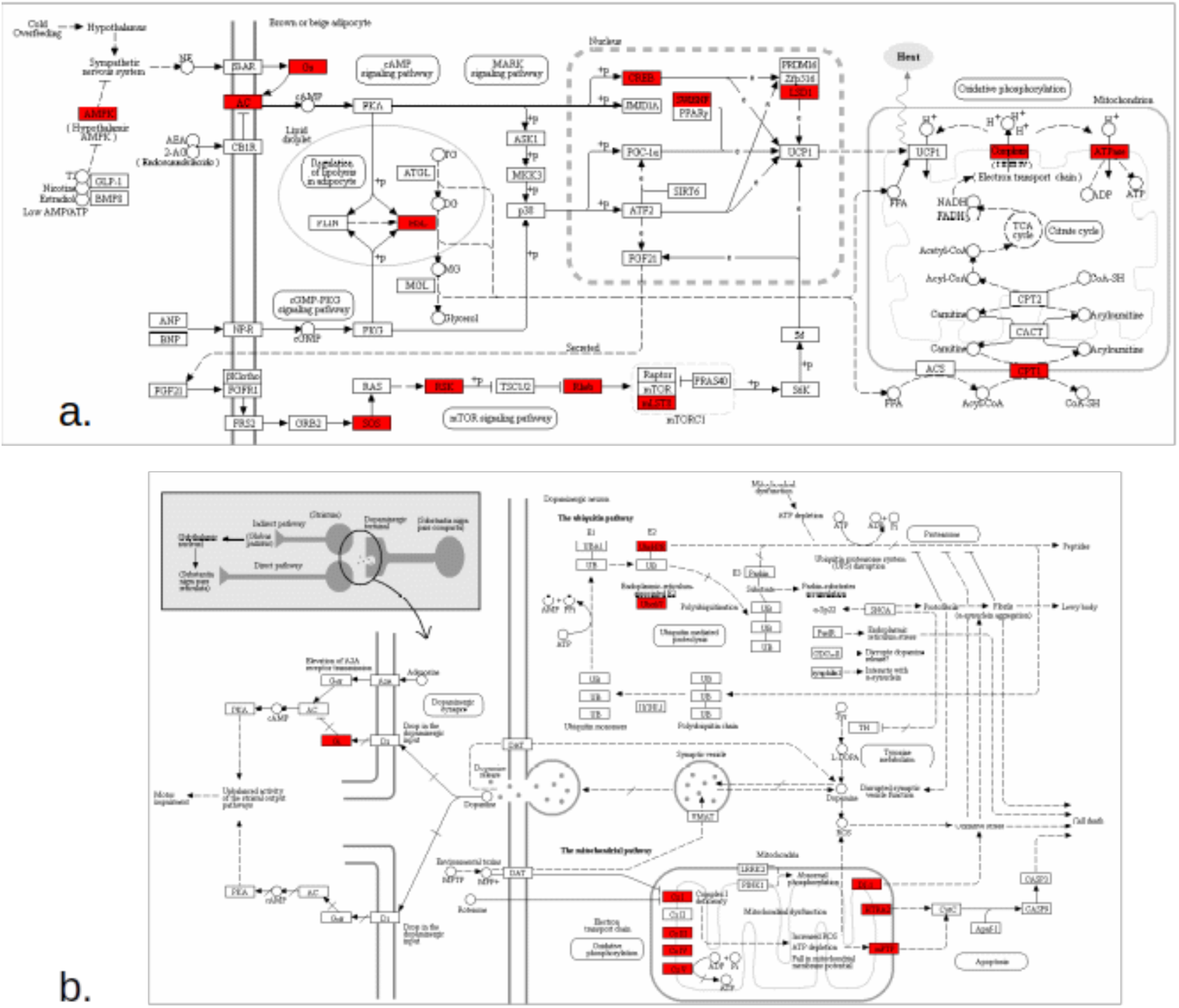
(Ridiculogram for; (a) Thermogenesis pathway rendered using pathview program integrated in ShinyGO, KEGG graph shows involvement of mitochondria in oxidative phosphorylation; (b) Parkinson’s Disease pathway rendered using pathview program integrated in ShinyGO; KEGG graph shows involvement of mitochondria’s inner compartment)

### 3.3 Mechanistic Analysis

Our analysis on correlated DE lncRNA and DE protein-coding mRNA pairs that were derived from progressor class-wise testing show that mRNA-PBX2 and lncRNA-AL662884.1 share common start bp on chromosome 6 (Table 2). In the same pair set (i.e. from progressor classes) Hypoxia-Inducible Lipid Droplet Associated (HILPDA) was mapped in close proximity (start bp within 1000 bp range) to AC010655.4 and AC010655.2 on chromosome 7. But in the case of correlated DE lncRNA and DE protein-coding mRNA pairs that were derived from non progressor class-wise testing no close elements were found (Table 2).

**Table 2.** (Chromosomal Location of the closely situated correlated DE lncRNA and DE protein-coding mRNA derived from progressor class-wise testing, i.e. “Mtb Infected” Vs “ESAT/Ag85 Stimulated”)

| mRNA Symbol | lncRNA Symbol | Chromosome | Distance |
| --- | --- | --- | --- |
| PNN | AL132639.3 | 14 | 298 |
| HILPDA | AC010655.4 | 7 | 9 |
| PBX2 | AL662884.1 | 6 | 0 |
| HILPDA | AC010655.2 | 7 | -88 |
| MNT | AC006435.1 | 17 | -774 |

The chromosomal distribution of the DE lncRNA derived from progressor class-wise testing was significantly different in comparison to the chromosomal distribution of the DE lncRNA from non-progressor class-wise testing (Fig. 7(a)). In progressor class-wise testing most of the DE lncRNA were expressed from chromosome 19, whereas in the case of non-progressor class-wise testing chromosome 1 being the largest chromosome showed most expression of DE lncRNA (Fig. 7(b)).

**Figure 7.**
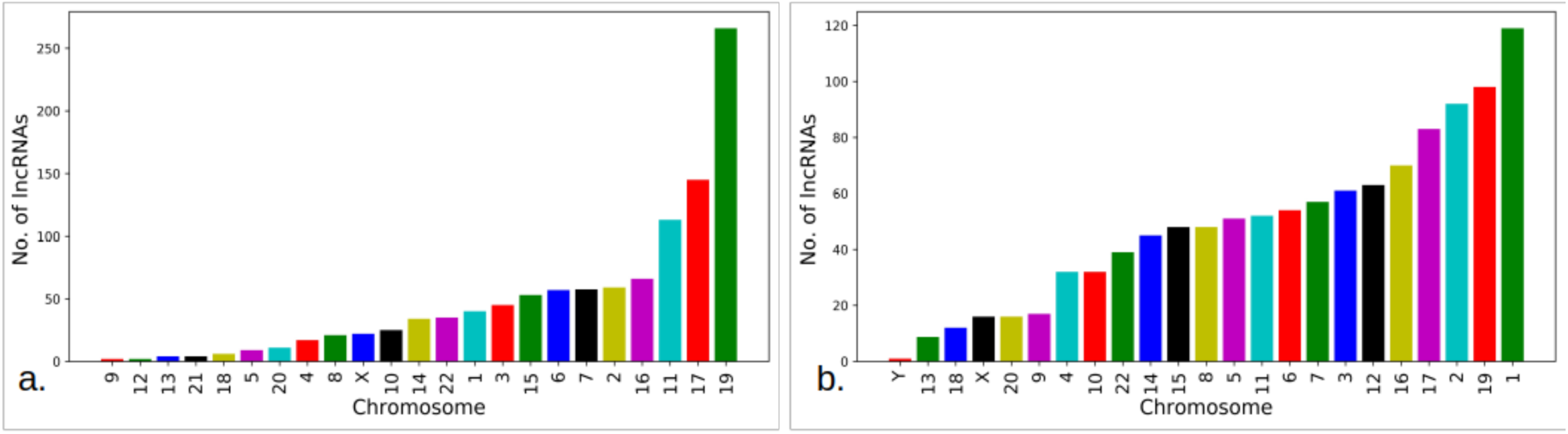
((a) DE lncRNA distribution across the human genome; Chromosome 19 being the smaller chromosome holds a higher number of DE lncRNA as compared to the chromosome 1,2 and 3; (b) DE lncRNA distribution across the human genome for non-progressors class-wise testing; Chromosome 1 is the largest chromosome holds a higher number of DE lncRNA)

**Figure 8.**
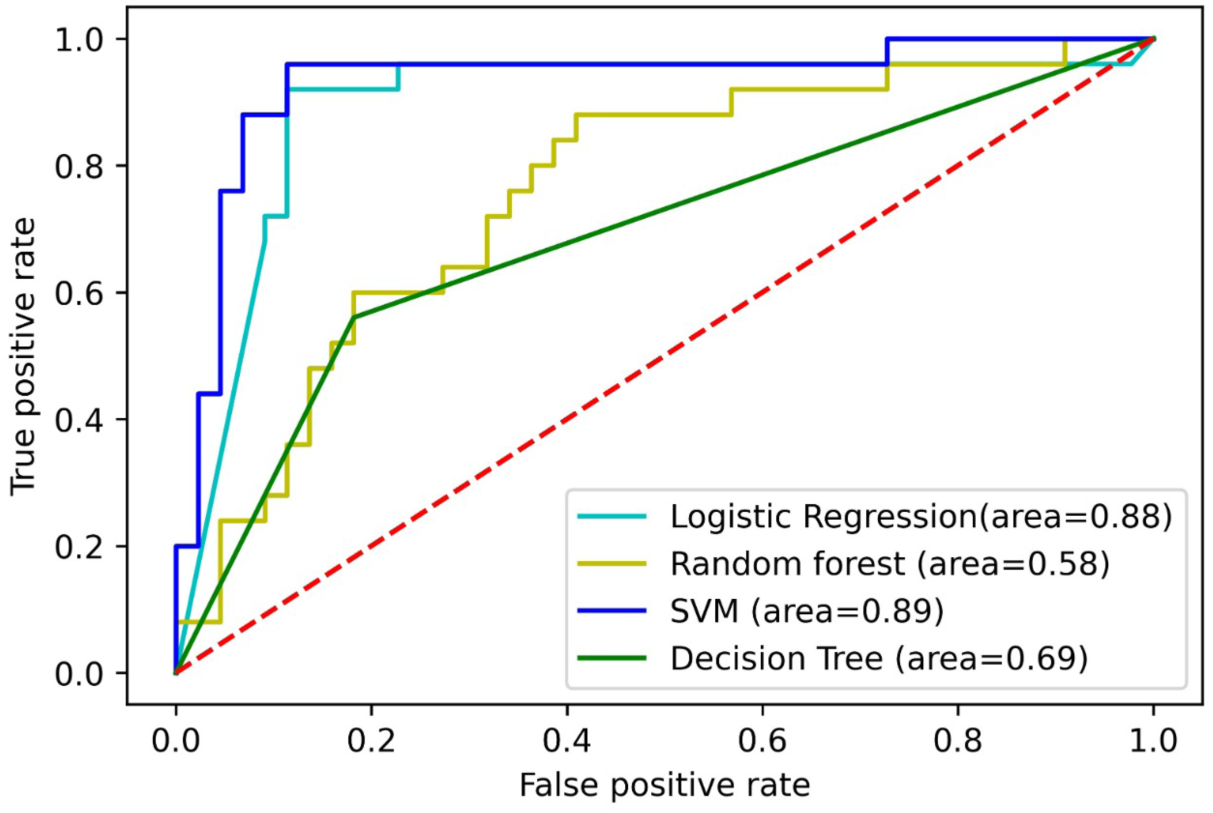
(Receiver Operating Characteristic curve of the models; SVM gave the highest accuracy of 89.85% with 84.61% sensitivity. In terms of specificity the LR outperformed the SVM with a specificity of 94.87%)

### 3.4 Transcription Factor Binding

After the transcription factor binding site prediction on the HILPDA-AC010655.4 and AC010655.2 trio we found 10 (ATOH7, BARX2, GSC2, NFIC, NFKB2, NFYA, NHLHI, SOX10, VAX1) common transcription factors among them. All of the DE lncRNA in class-wise testing showed insignificant (i.e. 0) expression potential.

### 3.5 Subcellular localization

Subcellular localization analysis revealed the location of 175 DE lncRNAs from the set of 529 DE lncRNAs in Mtb infected progressor replicates. Out of 175 DE lncRNAs, 125 DE lncRNA were enriched in the nucleus with uniform expression status (upregulation/downregulation) distribution i.e. 63 as upregulated and 62 as downregulated. 51 DE lncRNA from Mtb infected progressor replicates were enriched in cytosol, in which 40 DE lncRNAs were upregulated in comparison to only 11 being downregulated. Interestingly our finding enriched lncRNA-HIF1A-AS2 in the cytosol which is an anti-sense to HIF1A. The subcellular localization of the DE lncRNAs from non-progressive class-wise testing also showed uniform distribution both in nucleus and cytosol. The major upregulation of DE lncRNAs of Mtb infected progressor replicates clearly shows the evidence for cytosolic activity. As result of this non random enrichment we trained the model along with the DE lncRNAs of Mtb infected progressor replicates.

### 3.6 Classifier Predictions

SVM and LR classifiers were found to be independently useful on the basis of model performance indicators (Table 3), While SVM had a highest F1 score (89%) and sensitivity (84.61%) for predicting TB progressors, LR had the best specificity (93.02%) and overall second best accuracy (86.95%). Therefore, the LR model can be used for decisions to aggressively treat with a low false-positive rate whereas SVM can be used to triage for treatment escalation.

**Table 3.** Model Performance Indicators for Prediction of TB Progression.

|  | Logistic Regression | SVM Classifier | Random Forest | Decision Tree |
| --- | --- | --- | --- | --- |
| Accuracy | 86.95% | 89.85% | 68.11% | 72.46% |
| Sensitivity | 76.66% | 84.61% | 61.53% | 63.63% |
| Specificity | 94.87% | 93.02% | 69.64% | 76.59% |
| F 1 Score | 0.87 | 0.90 | 0.65 | 0.72 |
| ROC-AUC | 0.88 | 0.89 | 0.58 | 0.69 |

## Discussion

Our comparative study between the class-wise testing (i.e. “Mtb Infected” Vs “ESAT/Ag85 Stimulated”) in TB progressor replicates and non-progressor replicates showed a clear distinction. Upon validating with KEGG pathway enrichment analysis, 34 of the DE lncRNA from progressor class-wise testing, were enriched with Parkinson’s Disease pathway and Thermosgeneis pathway combined. Interestingly in both the KEGG pathways DE lncRNA sets were enriched within the lumen of Mitochondria whereas non-progressive class-wise testing showed no such shreds of evidence. Concrete evidence for many mitochondria-associated lncRNAs in the regulation of mitochondrial bioenergetics and cross-talk with nuclei has already existed in literature [38]. Also, during the cellular hypoxia mitochondria is the first organelle to be reprogrammed.. This non-random enrichment of mitochondrial compartments in the KEGG pathways directly points towards the involvement of mitochondria in TB progression from latent to active stage.

Our experimental design also enriched NEAT1 as an upregulated DE protein coding mRNA in the class-wise testing of progressive replicates. Role of NEAT1 is widely studied for its paraspeckle formation and paraspeckle dependent cell differentiation [39]. Upregulated levels of NEAT1 could manipulate the cytokine expression through the JNK/ERK MAPK signalling pathway in Macrophages [40]. It has also been reported as a potential biomarker in the previous studies [40]. Our study proposes 118 downregulated DE protein-coding mRNAs which are highly correlated with the low expressions of NEAT1 in class-wise testing if progressive replicates. Since NEAT1 is involved in nuclear retention of A-I mRNAs, therefore, it might be responsible architecturally blocking the expression of 118 DE protein-coding mRNAs in the nucleus. This could be backed up by the gene ontologies which exclusively enriched nucleus and nuclear functions as biological processes and cellular compartments (Table 1).

After the formation of hypoxic granuloma by the host immune system, T cells surround the infected macrophages to contain the infection. CD8+ cells being cytotoxic digs deep inside the granuloma to kill the infected macrophages [9]. For effective cytotoxic functions, T cells undergo hypoxic reprogramming which results in the secretion of hypoxic genes such as HILPDA and HI1FA. Our class-wise testing on non-progressors samples shows the overexpression of lncRNA-HIF1A-AS2 in the cytosol as well, which is antisense to HIF1A. This clearly shows that T cells undergo hypoxic stress even during latent stage. Interestingly there was no expression of HILPDA which is mainly responsible for lipid accumulation during the hypoxic condition. To withstand the hypoxic environment HIF1A is induced by the T cells. Our finding enriched lncRNA-HIF1A-AS2 in the cytosol which is an anti-sense to HIF1A. Coding potential of lncRNA-HIF1A-AS2 was very low, which clearly indicated it’s non-protein coding nature. Thus, expression and localization of lncRNA-HIF1A-AS2 in the cytosol are clearly due to the regulation of cytosolic functions of lncRNAs such as RNAi. In our expression set, the levels of HIF1A fail to pass the cut-off criteria of P-value but were majorly upregulated across the samples. Therefore, it is very much evident from our data that lncRNA-HIF1A-AS2 could mask the translation of HIF1A in the cytosol through RNAi.

We also report novel lncRNA-AC010655 (AC010655.2 and AC010655.4) for their consensus binding to HILPDA linked transcription factors in the nucleus. HILPDA is responsible for lipid accumulation as the cell slowly changes its metabolism towards low oxygen environments. Mtb has been known to adapt to a fatty-acid-rich environment of the cell [41]. It has also been experimentally proved that cellular organelles which store host lipids, are actively manipulated by Mtb during active TB [42]. This change is reflected in mitochondrial pathways as cells change their metabolism towards fatty acids, which are actively enriched in our data set of DE protein-coding mRNAs. Our results also show that non-progressors replicates of TB show the expression of HILPDA as upregulated protein-coding mRNA. Therefore, we report HILPDA as one of the major biomarkers of TB that is expressed both in latent and active stage.

In order to exploit these mechanistic insights, we chose to construct machine learning models that leverage the exclusive presence of a cytosolic signal in lncRNAs in combination with the differentially expressed mRNAs. Mechanistically enhanced machine learning models have not been applied to predict TB progression to the best of our knowledge and may avoid the pitfalls of black-box predictions which may not be actionable in the real-world settings. Our models also revealed an interesting conflict between the decision to choose between the overall best model and the potential to pivot clinical decision making and therapeutic implications. SVM, though the best performing model based upon F1 score was more sensitive, but not high enough to change decisions. On the other hand, LR classifier had a high specificity (94.87%) but a marginally lower F1 score. We propose the use of LR classifier, which has a low false-positive rate for informing decisions to escalate treatment in predicted progressors. On the other hand, SVM may be useful for triaging predicted progressors for treatment escalation as it has higher sensitivity. This study has several limitations. The mechanistic insights are derived from the correlation between DE lncRNA and DE protein-coding mRNAs and using DE protein-coding mRNA enrichment in pathways as a surrogate for DE lncRNA. We have attempted to mitigate some of the false associations by restricting ourselves to the DE lncRNAs that are present in cis with the correlated DE protein-coding mRNA. However, this may have led to missing out on the transacting DE lncRNAs. We also enriched for this effect by investigating the transcription factor binding sites that are common to the DE lncRNA-DE protein-coding mRNA pairs, thus increasing the probability of a functional interaction, which can be confirmed only through wet-lab experiments. Nonetheless, our study revealed a strong signal for the modulation of the hypoxic response pathway which is evident in Mtb infection but not in antigenic stimulation, thus indicating the hijacking of this machinery by Mtb specifically through cytoplasmically enriched DE lncRNAs. Finally, our prediction models are highly specific, but these can guide decisions in persons who may already be suspect for TB progression (confirmation) as opposed to a sensitive screen.

## Conclusion

Our study concludes that a non-random enrichment of DE lncRNAs in the cytoplasm, specifically those associated with the hypoxic response pathways dictates the progression of Mtb from latent to an active state. Our analyses indicate the mechanisms of such regulation and its potential to predict progression using machine learning models.

## Acknowledgements

This work was partly supported by the Wellcome Trust/DBT India Alliance Fellowship IA/CPHE/14/1/501504 awarded to Tavpritesh Sethi and the Tavpritesh Sethi also acknowledges support from the DBT Project BT/PR/34245/AI/133/9/2019 and the Center for Artificial Intelligence at IIIT-Delhi. Tavpritesh Sethi also acknowledges the valuable inputs provided by Dr Mitali Mukerji (CSIR-IGIB) and Dr Gaurav Ahuja (Computational Biology Department, IIIT-Delhi).

